# A Novel Computational Framework to Predict the Impact of a Point Mutation on PDZ Domain Classification

**DOI:** 10.1101/244251

**Authors:** Muhammad Moinuddin, Wasim Aftab, Adnan Memic

## Abstract

PDZ domains represent one of the most common protein homology regions playing key roles in several diseases. Point mutations (PM) in amino acid primary sequence of PDZ domains can alter domain functions by affecting for example, downstream phosphorylation, a pivotal process in biology. Our goal in this present study was to introduce a novel approach to investigate how point mutations within the Class 1, Class 2 and Class 1–2 PDZ domains could affect the changes in binding with their partner ligands and hence affect their classification. We focused on features in PDZ domains of various species including human, rat and mouse. However, our work represents a generic computational framework that could be used to analyze PM in any given PDZ sequence. We have adopted two different approaches to investigate the impact of PM. In the first approach, we have developed a statistical model using bigram frequencies of amino acids and employed six different similarity measures to contrast the bigram frequency distribution of a wild type sequence relevant to its point mutants. In the next approach, we developed a statistical method that incorporates the impact of bigram frequency history associated with each mutational site that we call history weighted conditional change in probabilities. In this PM study, we observed that the history weighted method performs best when compared to all other methods studied in terms of picking up sites in PDZ domain where a PM could flip the class. We anticipate that this method will present a step forward towards computational techniques unveiling PDZ domain point mutants that largely affect the protein-ligand binding, specificity and affinity. We hope that this and future studies could aid therapy in which PDZ mutations have been implicated as the main disease drivers such as the Usher syndrome.

## Introduction

PDZs are structural domains contained in many otherwise unrelated proteins. They are made up of approximately 90 amino acids and represent one of the most common protein domains found in the human genome [1–6]. PDZ domains are mainly recognized for their mediator role in the assembly of receptors at the cellular membrane interface [7–9]. For membrane scaffolding proteins these processes are often governed by PDZ domain binding of C-termini of their ligand protein partner, more specifically the last four to six amino acid residues [10–12]. Although, several PDZ domain classifications have been proposed, the three most dominant recognition patterns are *X*-*S*/*T*-*X*-Φ for Class I PDZ domains, *X*-Φ-*X*-Φ for Class II PDZ domains and Class I-II PDZ domains prefer the sequence of *D/E-X-*Φ (where Φ denotes an hydrophobic and *X* any amino acid)[13, 14]. NMR and crystallography structures of PDZ domains show that most of them share a similar three dimensional (3D) globular module [15]. In general, PDZ domains are composed of two α helices and six β strands. Several study implicated the pivotal role of PDZ domains in numerous diseases, mainly mediated by the PDZ domain interactions they contain [16–18].

Like many before, Tonikian et al. [19] showed that the most C-terminal ligand positions (position 0) and −1, and −2 positions as one moves towards the N-terminus are universally recognized by all PDZ domains in their dataset. However, they have demonstrated that up to eight distinct specificities for position 0 can exist. This strongly suggests that PDZ domains display significant promiscuity in their binding preferences both in terms of specificity and affinity. The primary driver for PDZ domain classification is the interaction between the domain residues at position within αβ1 of helix αβ and the 0 and −2 residue of the –COOH termini of a ligand [20]. In another report, by incorporating T7 phage display experiments Sharma, Sudhir C. et al.[6] have shown specification and bias towards amino acid (AA) residues on certain positions on the C terminal ligands that bind to Class I PDZ domains. For example, there is strong preference for either arginine or lysine at ligand positionP_-4_, and biased towards arginine at P_-1_.

Although, PDZ domains canonically interact with the proteins by binding to the C-terminal peptides, they are also capable of interacting with internal peptide sequences. One such well characterized interaction is between the PDZ domain of syntrophin and neuronal nitric oxide synthase (nNOS) [21, 22], where PDZ protein motif of N-terminus of nNOS, interacts with similar motifs in postsynaptic density (PSD)-95 protein and PSD-93.

Since PDZ domains are key players behind several crucial cellular processes for example, influencing synaptic protein composition and structure [2], mediating GIPC/Synectin associated interactions [5], enhancement of biomolecular associations in halogenated protein bindings [4], organizing binding of nNOS to skeletal muscle syntrophin [23] etc., hence it is not difficult to anticipate that point mutations (PMs) in PDZ domains can significantly impact various biological processes. Several other past and present works in this filed also suggests that PMs in PDZ domains can play pivotal roles in influencing promiscuity by altering the domain’s binding specificity [19, 24]. It is important to mention that in an experimental study reported by Tonikian et al. [19], specificity profiling of mutants of a model PDZ domain were reported and had been shown that ligand binding specificity is unaffected by most mutations because the C-terminal peptides were recognized by all mutants. However, it had also been suggested that binding specificity could change under mutational pressure.

In recent past, studies have shown that, mutations in the PDZ domain of harmonin has caused type 1C Usher syndrome [25–27], which is a genetic disorder resulting in visual impairment and hearing loss. This was one of the first reports that link the notion of critical mutations in PDZ domains and a disease state. Subsequent reports also described mutations in PDZ domain that could cause Dejerine-Sottas neuropathy, a lethal demyelinating disease causing peripheral neuropathy [28, 29].

When a group of PDZ domains exhibit a strong interaction preference towards a particular protein then they often share overlapping binding motifs [1, 30]. Challenges remain to engineer peptide ligands that only interact with a particular PDZ domain and specifically inhibit its activity. Vouilleme et al. [30] have investigated this issue on a group of five PDZ domains which were earlier found to be binding partners of the cystic fibrosis (CF) transmembrane conductance regulator (CFTR). They performed amino acid substitutions on 80 best HumLib peptides that interacts with their test PDZ domains. Their results confirmed that all of their test PDZ domains have differential preferences for the amino acids at P^0^. Similarly, Stiffler et al. [31] conducted an experimental study on 157 mouse PDZ domains aimed at quantifying their binding selectivity relative to 217 genome-encoded C-terminal peptide fragments. Their study demonstrated that sequence variations within the binding pocket of a given PDZ domains is often most correlated with differential binding preferences with the interacting ligands. In other words, a point mutation in the binding site makes a PDZ domain more prone towards a class flip i.e. a different kind of peptide ligand class.

In addition to these experimental studies many computational methods have also been developed to study PDZ domains [3, 14, 32]. However, these studies were mainly focused in designing features that could yield lowest possible false discovery in PDZ domain classification. To the best of our knowledge, so far no generic computational method exist that aid in pinpointing mutational sites in PDZ domain beyond the binding pocket that concurrently quantify the probability of misclassifications. These PM causing a classification flip could also be representative of a structural domain misfolding and could be relevant to disease study [25–28, 33]. Inspired by the literature in this field, we wanted to assess if similar principles can be applied to PDZ domain sequences in contrast to the previous studies that focused mainly on engineering peptides [30, 34]. Here, we developed a point mutation algorithm to mutate positions in PDZ domains and introduced a novel approach to study the impact of PM in the classification of PDZ domains. We have developed two statistical models for the PM study that are based on bigram frequencies of AAs and global history (defined later in Methods) of PDZ sequences. These models are based on various similarity measures (defined later in Methods) among them the Jaccard’s similarity performs best picking up significant mutation sites that we define as PM that could change ligand binding affinity and hence could possibly trigger a class flip [19]. Previous reports have identified critical sites in PDZ domains by grouping amino acids in seven groups based on their side chain volume and polarity [35], but our aim here is to identify critical PM formed by single amino acid substitution. In this context, the only significant work is of the McLaughlin RN, Jr. et al.[36]. However, their experimental study was limited to a specific PDZ domain, namely PSD-95. In contrast, our aim is to develop a generic computational method that can identify critical PM in any given PDZ domain based on the trends we observe from our dataset.

## Materials and Methods

In order investigate the impact of PM on PDZ domain classification, we extracted data from several sources including the work of Kalyoncu et al. [35]. Our data contained 45 Class I, 20 Class II and 21 Class I-II PDZ domains (See S1 Table). We used them to generate a bigram history for each class. However, in order develop and validate our generic mutant model we have used additional PDZ domains as wild type test sequences from protein database (PDB) [37] and also according to another previously published report [36]. The results and the impacts of PM on each of the test PDZ domains are discussed in detail in the Results and Discussions Section.

### Algorithm to generate mutants

The set ϗ = {A, C, D, E, F, G, H, I, K, L, M, N, P, Q, R, S, T, V, W, Y} has all possible AAs. We define a point mutation as a replacement of a single AA by another AA in the PDZ domain sequence.

Previous studies reported that PM can alter specificity of PDZ domains [19, 20, 38], however we were interested to investigate ‘which and how much’ from all possible PM effect the classification of PDZ domains. To do so we artificially introduced all possible pseudo-PM in our wild type test sequence, we denote it by P. We introduce three operators: mutation operator Җ, extraction operator Ѫ and length operator Ѧ.Җ(R, X) implies replace R by X, ∀X,R ∈ ϗ. Ѫ(ϗ(!X)) implies extract all AA from ϗ except X and convert into a string. Ѧ(S) returns the length of the sequence denoted by S.

Therefore, the PM algorithm can be written as follows,

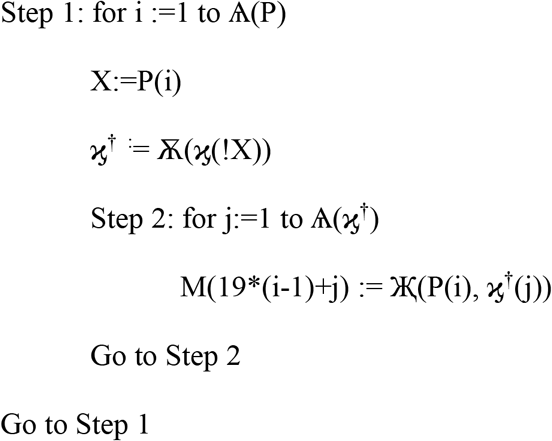

Fig 1 depicts the working and outcome of PM algorithm. Further, these new sequences were used as testing sequences during statistical analysis to check if any point mutation causes a class flip.

**Fig 1.**
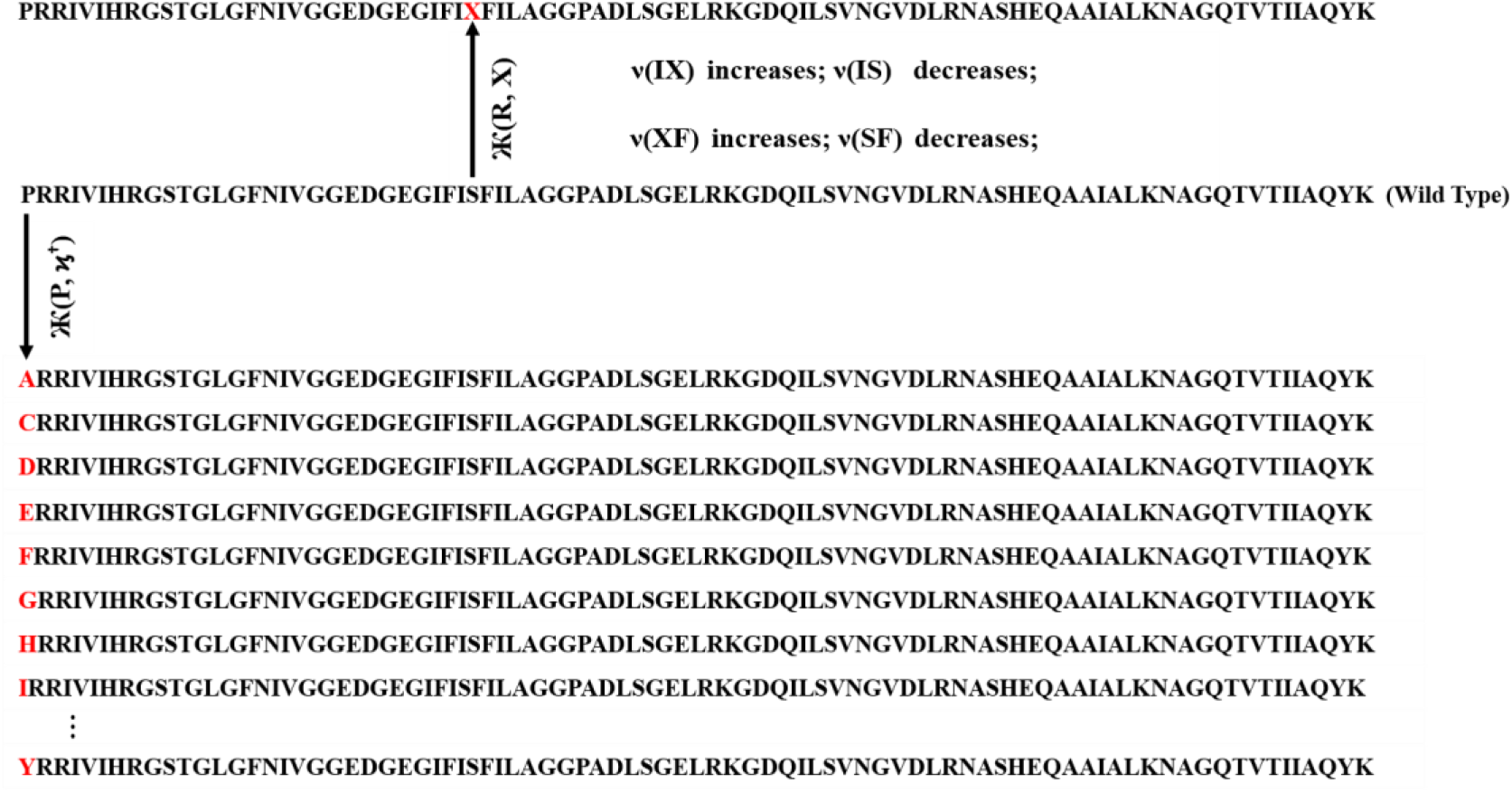
PM algorithm: Generating 19*L mutants. The mutants generated by mutating the first position only are shown. In addition, the effect of PM on BLFV is also shown.

### Feature Extraction from PDZ sequence

Features are key to our study. We have used the local and global bigram frequencies of amino acids as features in order extract features that best represent the training set.

### Bigram Local Frequency Vector (BLFV)

A protein sequence of length L, it has L-1 bigrams. For example, EITLERG sequence has the following bigrams: EI, IT, TL, LE, ER, and RG. We computed the bigram normalized frequency of amino acids in a PDZ sequence as follows,

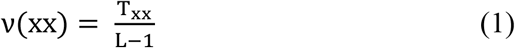

Where, x ∈ ϗ is an amino acid. This way we generate a probability distribution of bigrams from symbolic sequence of AA and *ν* is the BLFV. The reason of using normalized frequency of bigrams as features is that it converts a symbolic sequence of AA into a probability distribution and the idea of probability distribution is crucial in our study, which will be evident in subsection of the following section where we use normalized frequency of bigrams to investigate the impact of PMs in PDZ domains.

### Bigram Global History Matrix (BGHM)

In order to compute BGHM we first aligned our dataset separately for Class 1, II and I-II PDZ domains using CLUSTALX 2.0 [39]. The concept of BGHM is illustrated in Fig 2, where we compute the global frequencies of every existing bigram for a set of PDZ domains sequences. Since, we are dealing with three classes of PDZ; therefore, we have three BGHMs: G^I^, G^II^ and G^I-II^. Every Column in G^k^ (k=I, II, I-II) indicates normalized frequencies of all possible bigrams at that position in every PDZ domain. The conserved motifs across sequences are highlighted in logo diagrams 2(b)-(d) respectively for classes I, II and I-II PDZ domain using WebLogo [40]. The height of each stack in the logo diagram implicates conservation at that point in the sequence as measured by the bits present at that position. Similarly, the height of a symbol at each stack provides the relative frequency of the corresponding amino acid at that given position.

**Fig 2.**
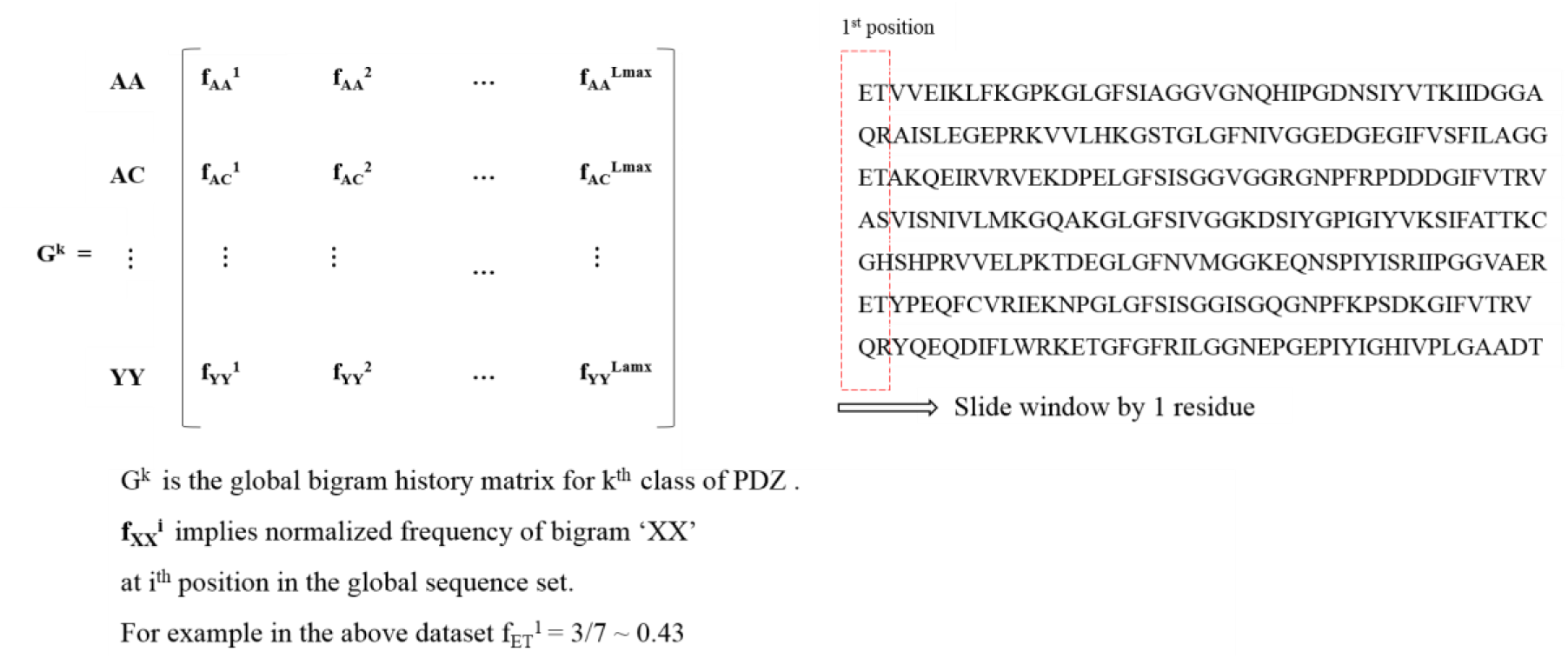
Concept of global bigram history matrix and sequence alignment. (a) Bigram Global History Matrix. Graphical representations of the patterns within a multiple sequence alignment. (b) Class I (c) Class II (d) Class I-II

In order to facilitate our computational methods we have chosen the maximum number of columns in G^k^ to be the same as that of length of the longest PDZ domain sequence for k^th^ class.

### Statistical Models for PM investigation

As we mentioned earlier that we have mapped each symbolic AA sequence into a bigram probability distribution BLFV. So we have a BLFV for wild type and 19*L BLFVs for all mutants that are generated using our PM algorithm. Our idea is to see which PM cause class changes, thereby observing how similar a BLFV is with its wild type after a PM.

#### Using only BLFV

We have used following similarity measures from the work of Sung-Hyuk Cha [41].

i. Euclidian Similarity
ii. Minkowski Similarity
iii. Tanimoto Similarity
iv. Jaccard Similarity
v. Czekanowski Similarity
vi. Cosine Similarity
vii. Motya Similarity

These definitions are presented in Table 1, where, P and Q are the BLFVs for wild and its mutant respectively, L represents the length of the PDZ domain sequence, p represents the degree of the root for a distance measure. However, it is important to note that since the symbolic equivalent of P and Q differs by only one amino acid; therefore; in the BLFVs of P and Q will differ at most in four positions due to the different bigrams. This idea is illustrated in Fig 1. Hence, we extracted only the bigram frequencies that differs in both wild and the mutant. The idea is explained with the help of following equation.

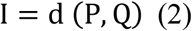

**Table 1.**
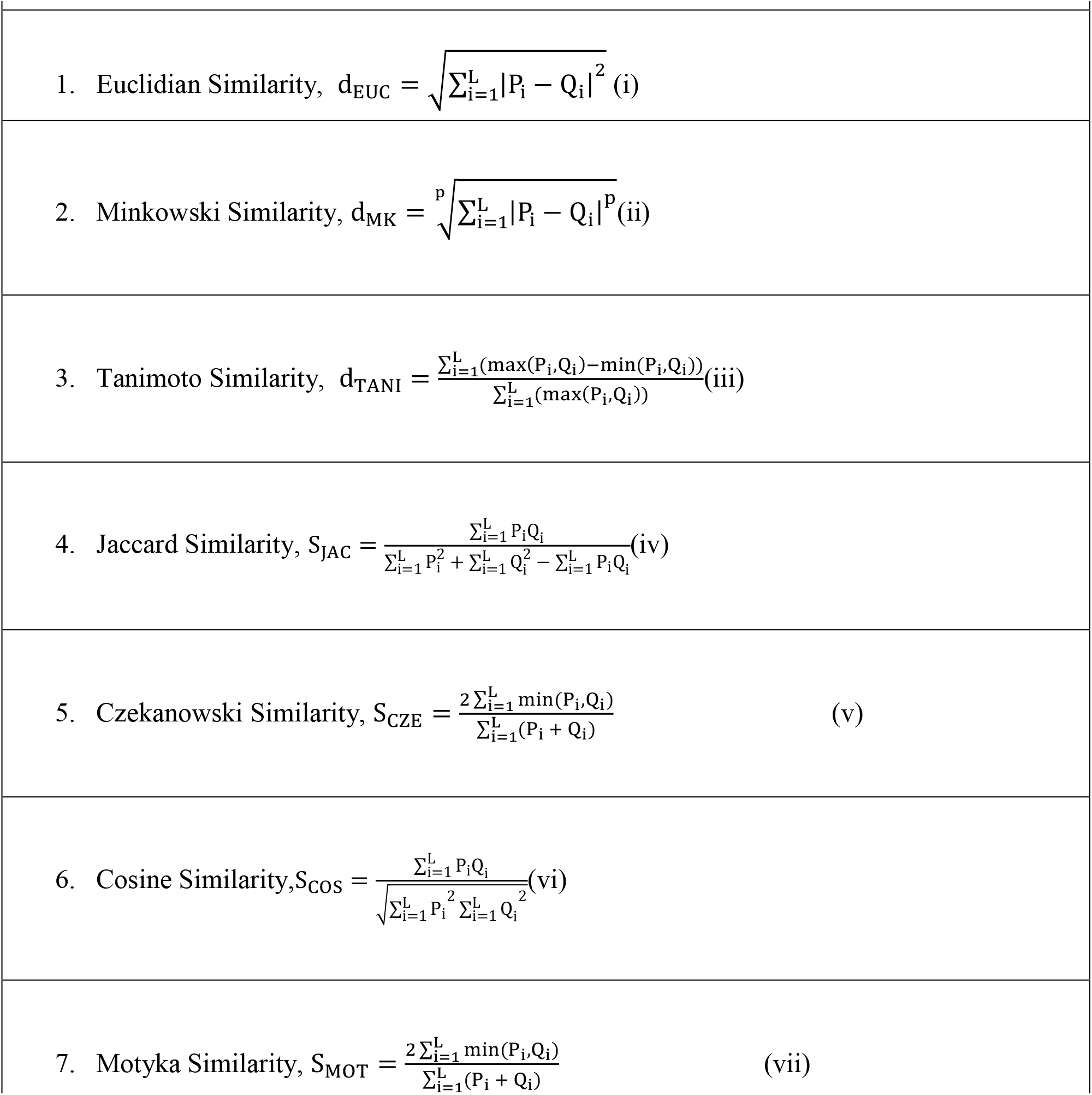
Definitions of Similarity/Distance measures.

I is an indicator variable that stores only those indices for which mutant differs from its wild type. Therefore, the Jaccard similarity index from the Eqn. (4) of Table 1 can be written as,

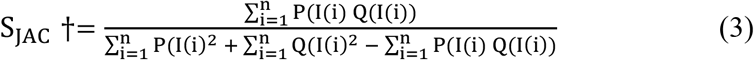

#### History weighted conditional change in bigram probability (HWCC) using BGHM

The BGHM G^k^ is computed as shown in Fig 2, where we have used a window of size two to extract global bigrams. The window starts sliding at 1^st^ residue position and moves towards right by one residue. The frequency of any bigram ‘XX’ in each window is computed using following equation.
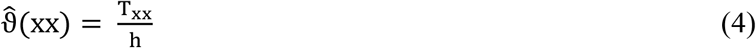

where, h is the window height or in other words, the number of bigrams in a window. T_xx_ is the frequency of XX in a window.

As discussed earlier that we have indicator variable I that store indices of only those places where wild type P differs from its mutant Q (See Eq. 2). In this section, we introduce an idea of history weighted conditional change in bigram probability. For every mutant we compute the total change in probability as follows,
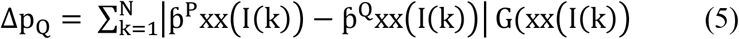

where, 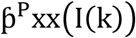 is the frequency of bigram XX at position I(k) in P, 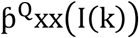 is the frequency of bigram XX at position I(k) in Q, N is the number of places P differs from Q, G(xx(I(k)) is the global frequency of bigram XX at position I(k). Finally, the decision variable for a mutant Q is computed using logsigmoid as,
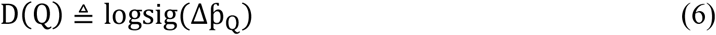

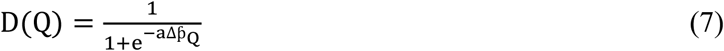

where, ‘a’ is sigmoid exponent. The reason for using log sigmoid function is just to amplify even small values in the result in 0–1 window. This way it shows up more distinctively in the heat map. The inclusion of history matrix provides additional weights to the local bigram frequencies and to the best of our knowledge, it has not been used before to investigate the similarity of two PDZ domains.

## Results and Discussion

### Class similarity after point mutation

We used third PDZ domain of PSD-95 protein (PDB: 1TP3) as test bed because effect of PM in it has already been studied experimentally [41]. Since we have introduced single point mutation in each position, therefore, the effective change is very tiny. Therefore, in order to capture that small change we needed more precise methods than one to one correlation between the mutant and the wild type, which are discussed as follows.

#### Using BLFV only

We have already mentioned that we use several similarity measures to capture the difference a mutation brings on the wild type. However, among the seven distance measures used, we found Jaccard and Cosine measure performed reasonably well while Jaccard being more sensitive in terms of accuracy with respect the published results in [36]. The heat maps are shown in Figs 3(a)-(g).

**Fig 3.**
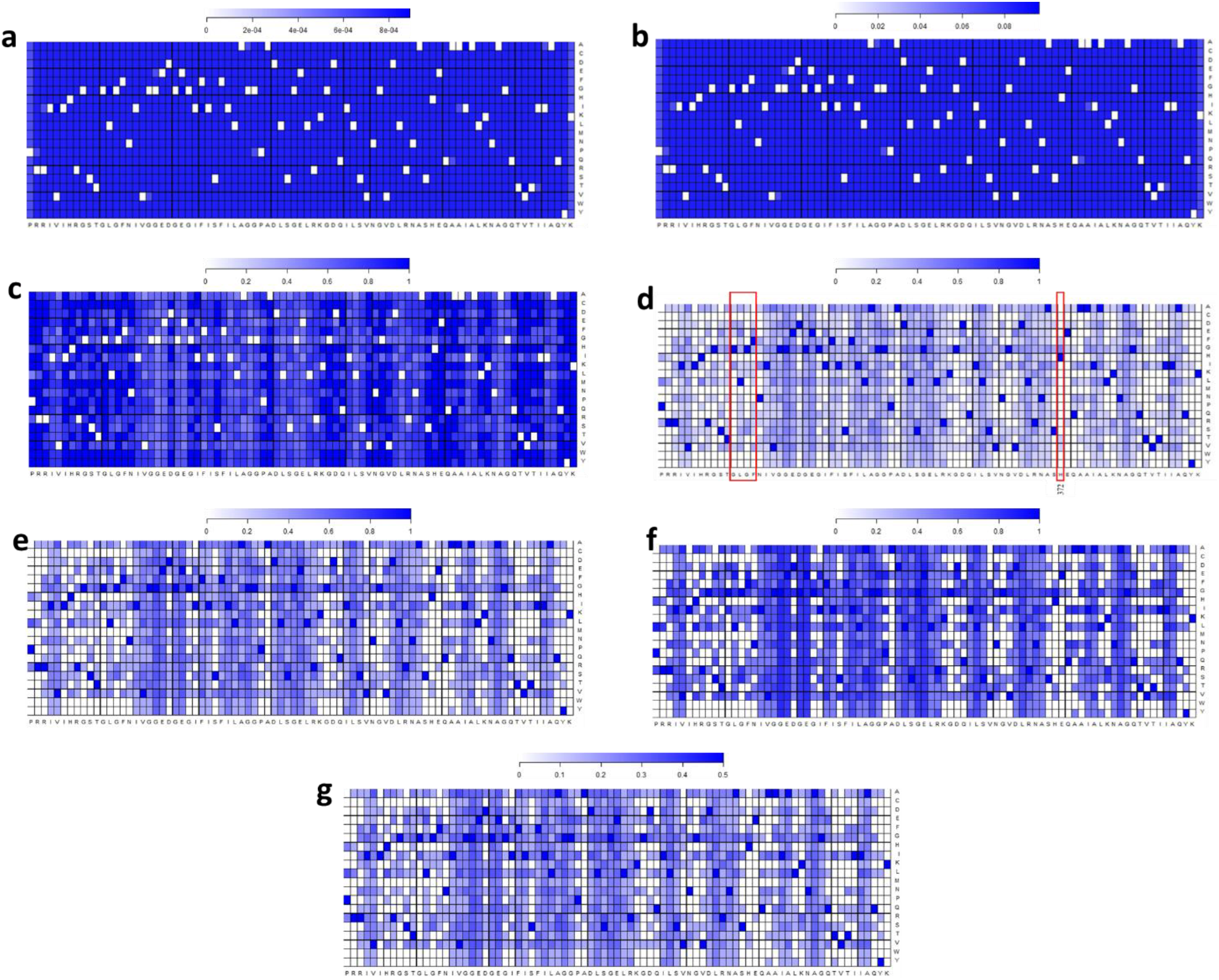
Mutation Heat Map for third PDZ domain of PSD-95 protein (PDB: 1TP3). Figures 3(a)-3(g) are computed based on following similarity measures between wild and mutant probability distribution respectively: Euclidean, Minowski, Tanimoto, Jaccard, Czekanowski, Cosine, Motya. In figures 3(a)-3(c), as one move towards blue shade in the color scale implicates that these positions are highly affected due to mutational pressure and whiter means probability of change in classification is less. In contrast, in figures 3(d)-3(g) whiter portions indicates that these positions are highly affected due to mutational pressure and more towards blue implies less probability in flipping class.

We used our library of PM sequences on Jaccard Similarity algorithm. Mutations on the binding pocket motif “GLGF” in 1TP3 is found to be altering the similarity among the mutants (Shown in Red box in Fig 3(d). Such flips were anticipated, as changes in the PDZ domain sequence changes the binding pocket and peptide interactions. The position 372 (Shown in Red box in Fig 3(d)) is found to be a highly affected by the mutation. The wild type residue Histidine (H) at that position forms hydrogen bonds (Shown in green dotted line in Fig 4) with alpha helix and ligand. We anticipate whenever a mutation is causing these bonds to be broken the classification similarity changes i.e. it also affects the ligand binding specificity of 1TP3 which is reflected in the Jaccard similarity scores at that position for each such mutation. In general, we found that most mutations away from binding site are not flipping the class, which is in line with the reports of Tonikian et al. [19], where they reported that PDZ domains are highly robust towards most point mutations. However, we found some PM away from the binding sites and in the loops are able to produce class flips. In general, loops are cysteine rich hence when a mutation alters the composition of AAs it could trigger an effect similar to allosteric effect, where perhaps PDZ domain is being regulated by the ligand binding at a site other than its active site. However, these hypotheses still require experimental verification. As mentioned earlier that Jaccard with BLFV performs comparatively better than others, but it produces some false mutations towards the beginning and the end of the sequence. Therefore, in order to reduce the false positives and to see if history of bigram frequencies has impact on the PM in PDZ domain we incorporated bigram history of amino acids into our study.

**Fig 4.**
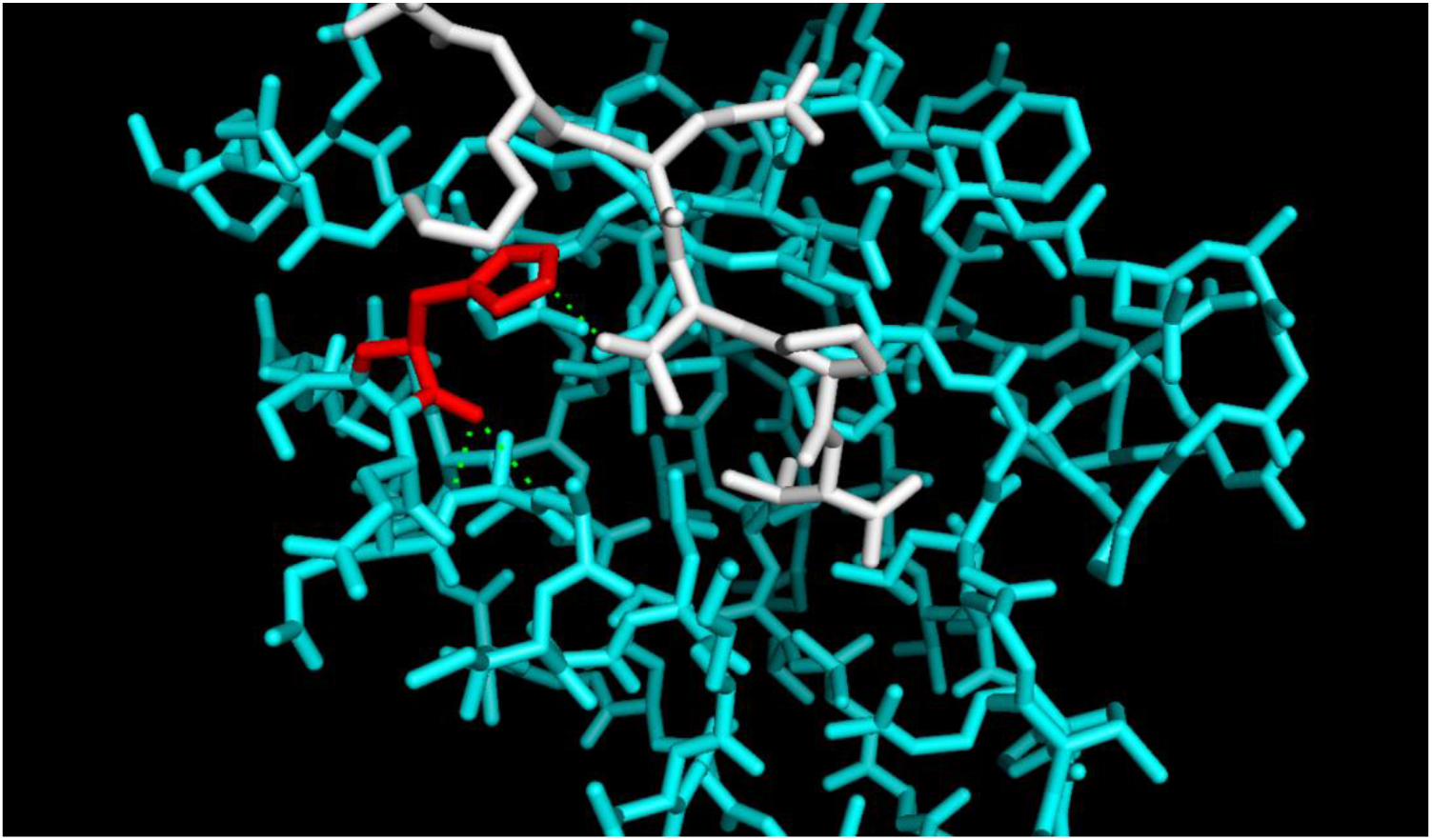
Stick representation of PDZ domain. Third PDZ domain (gray colored) of PSD-95 protein (PDB: 1TP3) complexed with KKETPV peptide ligand (blue colored); Histidine is shown in red color.

#### Using HWCC

The Jaccard’s similarity method appears to produce many false positives. To improve the statistical outcome and reduce possible false positives we introduce another method based on history of bigrams (See 3.3.2). The heat map using the history weighted conditional change in bigram probabilities are shown in Fig 5(a).When the heat map of Fig 5(a) is compared with that of Fig 3(d), we find that HWCC method performed much better in terms of picking up the mutations in the binding site motifs, while reducing the false positives.

**Fig 5.**
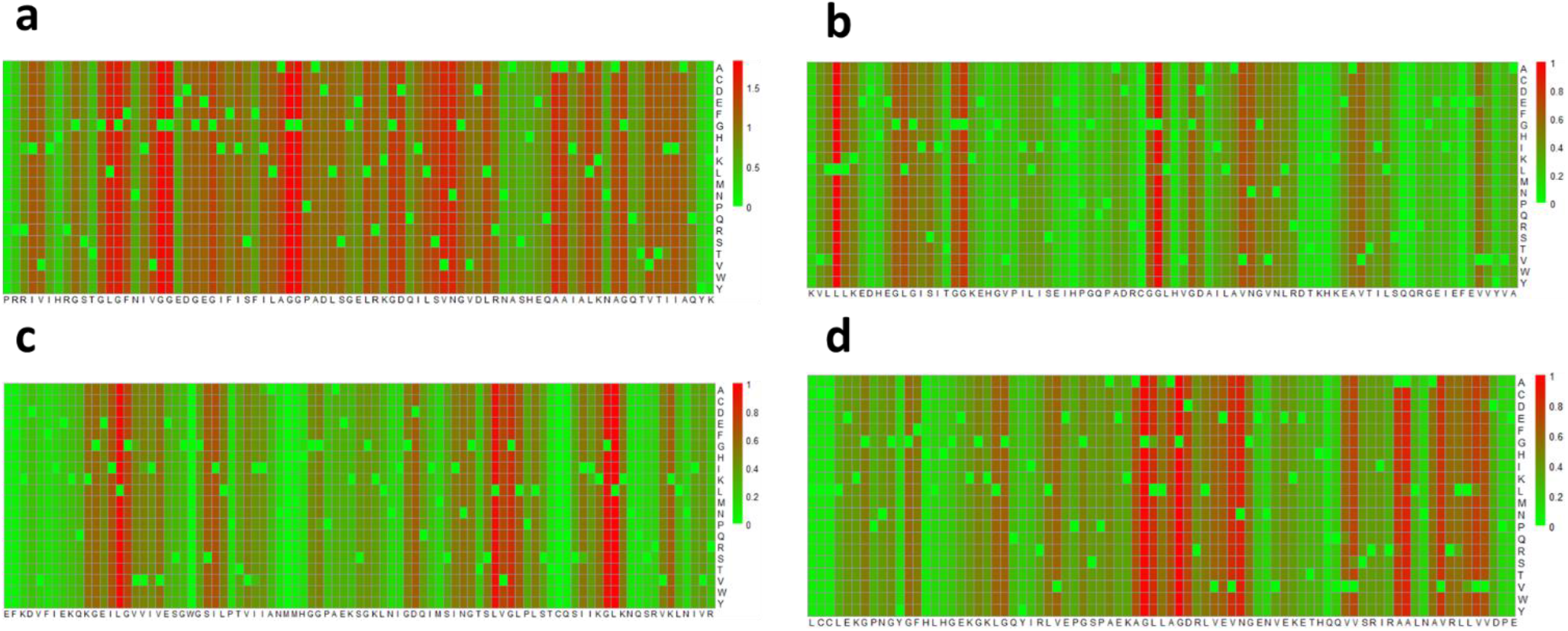
Heat maps using history weighted conditional change in bigram probability method. Mutation in motifs (shown in red) is prone to change classification. Critical motifs are shown for (a) third PDZ domain of PSD95 (PDB: 1TP3, Class 1), (b) CAL PDZ domain (PDB: 2LOB, Class 1), (c)PDZ domain of X11 proteins (PDB: 1U38, Class 2), (d) PDZ domain from NHERF1 (PDB: 2M0U, Class 1), (e) PDZ domain from PTP-BL (PDB: 1VJ6, Class 1–2), (f) PDZ domain from MAGI-1(PDB: 2I04, Class 1–2).

The HWCC method heat map in Fig 5(a) is much more in line with that of the work of McLaughlin Jr. et al. [36]. However, it still catches what appear to be a few false positives with respect to the experimental results [36], which also reflected in the Receiver operating characteristic (ROC) curve shown in Figure 7, where with AUC=0.72 we could not pick all experimentally determined mutations.

Therefore, our study aims to provide a generic computational approach to pick most of the important PM without having to go through lenghty experimental processes. In addition, this method highlights some possibly new critical PM sites, which might be of great importance in future disease studies. Hence, it opens new dimension of research to explore the biological significance of such PM sites through experiments.

However, it is important to mention our present method is highly dependent on the sequences in the database. For example, after multiple sequence alignments frequencies of all the highly redundant motifs increase by several fold while at the same time low frequency motifs frequencies can become significantly reduced.

In order to test robustness of HWCC method we downloaded some more PDZ sequences from PDB (PDB IDs: 2M0U, 2LOB, 1U38, 1VJ6, 2I04) [37]. The resulting heat maps are shown in Figs 5(b)-(f). We were able to identify key motifs as per our initial hypothesis. In all cases, we were able to detect mutations on and around the binding site that were responsible for changing the class conditional probability [42]. The surface image representations are in shown using Pymol in Figs 6(a)-(e).

**Fig 6.**
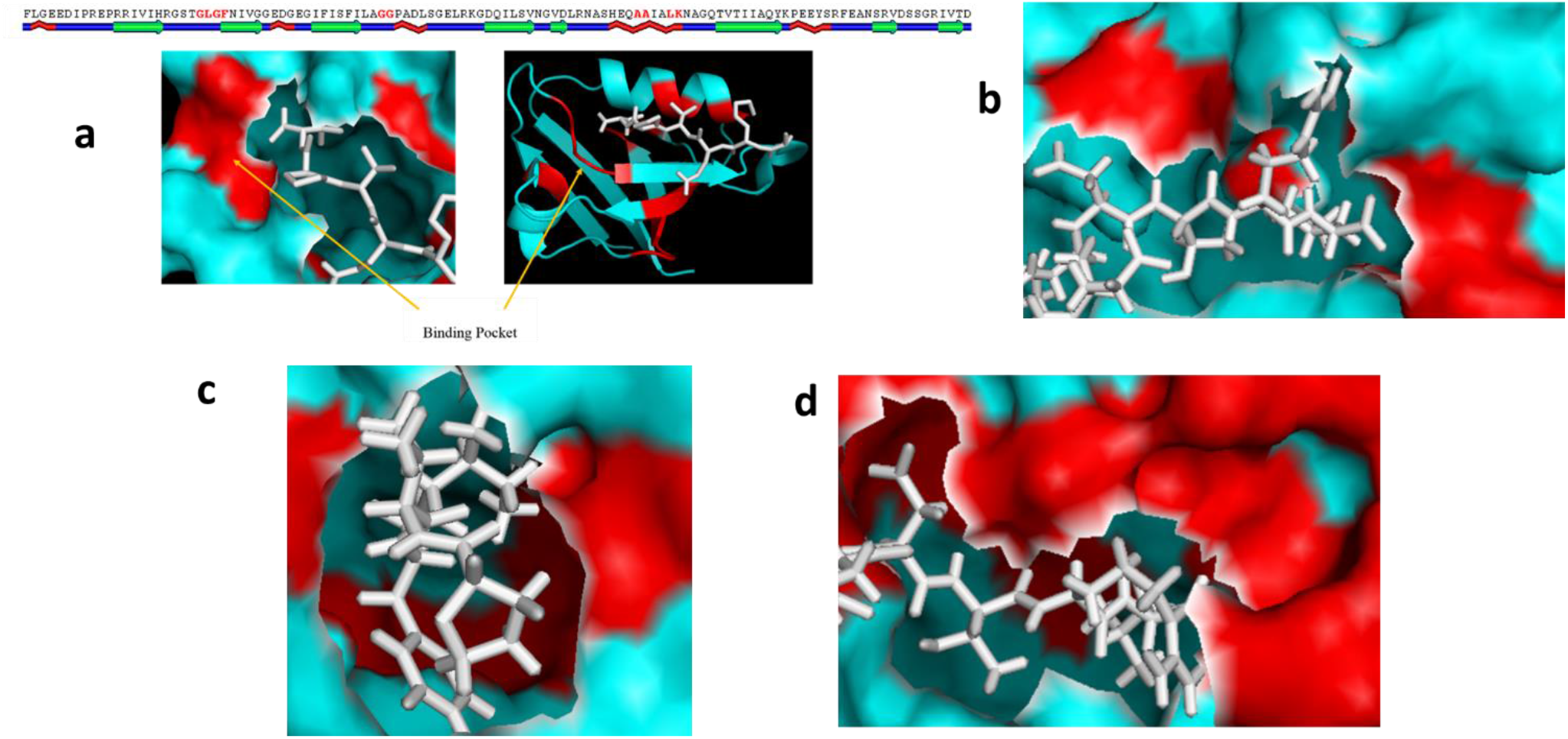
Graphical Visualization of Critical Motif Binding Sites of PDZ domains. (a) Critical motifs are shown in red for the third PDZ domain (cyan colored) of PSD-95 protein (PDB: 1TP3) complexed with KKETPV peptide ligand (white colored).Which are also highlighted in the tertiary ribbon view, a close up surface view of the binding site is also shown. (b) PDZ Domain of CAL (PDB: 2LOB), (c) PDZ domain from X11s/Mints Family Scaffold Proteins (PDB: 1U38), (d) first PDZ domain from NHERF1(PDB: 2M0U), (e)first PDZ domain form MAGI-1 (PDB: 2M0U)

**Fig 7.**
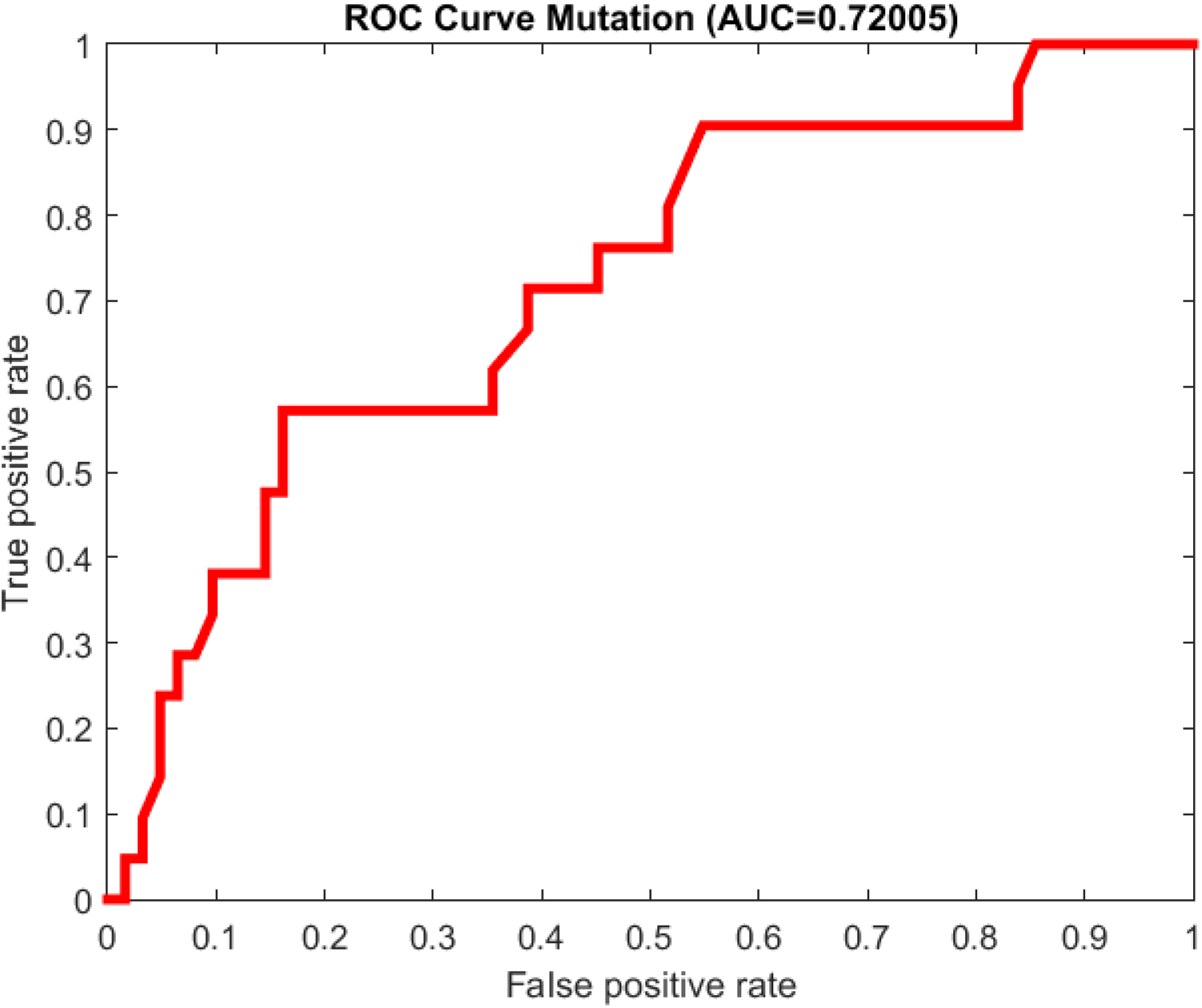
ROC plot to validate mutation on 3^rd^ PDZ domain of PSD-95[34] using HWCC method. This moderate ROC also supports heat map in Fig 5(a) where HWCC method did not catch all the reported mutation, however, it catches some other critical motifs (see next section) reported in literature[14].

### Critical Motifs

Several life-threatening diseases are caused by PM in linear motifs that mediate important interactions [43–45] and they are commonly refer to as Motif-Mediated Diseases (MMDs). For example, Usher’s Syndrome, has been shown to be the result of mutations in either PDZ domains present in the Harmonin protein or alternatively in the PDZ interaction motifs that interact with Harmonin such as the SANS protein [44]. After analyzing our heat maps in Fig 5 and its structural realizations in Fig 6, we noticed that certain motifs are more prone in flipping the class than others. Since we incorporated history weighted bigram frequencies of AA to identify these critical motifs, a PM in those motifs are historically more inclined in changing the classification of PDZ domains. In Fig 6(a), we show the critical motifs for third PDZ domain of PSD-95 using Polyview and Pymol [42, 46]. For the third PDZ domain of PSD-95 in Fig 6(a), our PM algorithm catches following critical motifs: “GLGF”, “GG”, “LK”, “AA”, “GD” etc., out of them “GLGF” and “AA” are binding site motifs hence very critical. The other motifs are also important as most of them are on the beta sheet or on the alpha helices. The bigram “GD” is reported to be critical by Kalyoncu et al. [35]. In Figs 6(b)-(d) we investigated and found additional critical motifs that include “DH”, “ED”, “EG”, “GI”, “GL”, “HE”, “IS”, “IT”, “KE”, “KV”, “LG”, “LK” and others. It is interesting to note that using HWCC method we were able capture the binding site motifs in all test cases and most motifs are found to be on the alpha helices/beta sheets, though some are caught on loops that connect these structural features.

It can be noted from Figs 2(b)-(d), where most of the discovered motifs are conserved throughout species, implying that any PM in these motifs could have a high correlation in altering binding specificity of PDZ domains and hence could possibly affect domain classification. It is important to note that these motifs were discovered after incorporating the bigram histories of amino acids. Therefore, these motifs are historically important too and more prone in flipping classification under mutational pressure than others.

Encouraged by the fact that our findings are strongly correlated by past results in this field [19, 36], and the significant role of PM in protein engineering in general and correlated to several life threatening diseases [43–45], we propose that several mutations found could be pivotal in understanding of disease states. However, further experimental studies are needed to in order to reveal biological significance of these PM and how large of an impact they play on PDZ-ligand interaction and in disease etiology.

## Conclusion

In this paper, we have introduced a model to study the impact of PM on PDZ domain classification. By incorporating history of bigram frequencies of amino acids in the conditional change in probability distributions we demonstrated that mutations not only in the binding pocket, but also far away from loops are possibly involved in canonical ligand recognition and able to affect PDZ domain classification. We also shown that mutations in the binding site motif highly destabilizes the domain structure. We attribute this to possible structural alterations that could ultimately affect the three dimensional protein fold. With accurate computational methods identifying PM motifs responsible for altered PDZ specificity, we hope that they can be used to optimize MMD research. Moreover, in cases where a strong relation exists between PM in PDZ domains and life threatening diseases this computational simulations could prove to be useful.

## Acknowledgement

We would like to thank Dr. Ahmed Balamesh for his insightful discussions and comments.

## Supporting Information

**S1 Table. List of PDZ domains used to construct history matrices.** Primary amino acid sequences of all PDZ domains used in the point mutation study detailing the organism, classification, PDZ domain ID, name and origin.

